# Metrics of Coral Microfragment Viability

**DOI:** 10.1101/2023.01.03.522625

**Authors:** Claire Lager, Riley Perry, Jonathan Daly, Christopher Page, Mindy Mizobe, Jessica Bouwmeester, Anthony N. Consiglio, Jake Carter, Matthew J. Powell-Palm, Mary Hagedorn

## Abstract

Coral reefs are being degraded at unprecedented rates and decisive intervention actions are urgently needed to help them. One such intervention in aid of reefs is coral cryopreservation. Although the cryopreservation of coral sperm and larvae has been achieved, preservation of coral fragments including both its tissue and skeleton, has not. The goal of this paper was to understand and assess the physiological stressors that might underlie coral fragment cryopreservation and the long-term consequences of these physiological exposures to continued growth. Therefore, we assessed small fragments (∼0.5 x0.5 mm^2^) from the Hawaiian coral, *Porites compressa*, examining: 1) the sensitivity of the fragments and their algal symbionts to chilling temperatures; 2) the sensitivity of the coral to complex cryoprotectants; 3) methods to safely remove the algal symbionts from the coral fragment for cryopreservation, given the two symbiotic partners may require different cryopreservation protocols; 4) continued growth over time of coral fragments once returned to running seawater after treatment exposures; and, 5) assessment of health and viability of microfragments after treatments examining the distribution of green fluorescent protein and fluorescent symbionts. Technological advances in cryo-technology promise to support successful coral fragment cryopreservation soon, and its success could help secure much of the genetic and biodiversity of reefs in the next decade.

## Introduction

Throughout the world coral reefs are being degraded at unprecedented rates. Locally, reefs are damaged by pollution, nutrients and sedimentation from outdated land-use, fishing and mining practices, and globally, increased greenhouse gases are warming and acidifying oceans, making corals more susceptible to stress, bleaching and newly emerging diseases (Eakin et al. 2019; França et al. 2020; Frölicher et al. 2018; Hughes et al. 2017; Muller et al. 2020).

Although well-managed marine parks are afforded some protection, our reefs remain vulnerable to disease, natural disasters and human impacts. Because these threats do not respect geo-political boundaries, we must resolve to act globally to implement effective conservation strategies. Although *in situ* conservation strategies, such as marine protected areas and coral restoration, may help mitigate local threats to coral reefs, the global effects of climate change will continue to erode these important areas causing steady declines in coral population numbers.

The coupling of climate change and anthropogenic stressors has caused a widespread and well-recognized reef crisis (Bellwood et al. 2004; Madin & Madin 2015). New modeling data suggest that the threat to tropical coral reefs may be challenging even with the most optimistic assumptions of coral reef refugia, adaptation and potential for restoration with near total reef loss expected by mid-century (Dixon et al. 2022; Kalmus et al. 2022). As part of these stressors, ocean warming is increasing the frequency of bleaching events around the world (Hughes et al. 2018), negatively impacting coral reproduction (Hagedorn et al. 2016; Ward et al. 2002). Without robust reproduction on reefs, the potential for adaptations to warmer waters is reduced (van Oppen et al. 2015). Because the next 10 to 20 years will be so perilous in terms of loss of species for our oceans, we need innovative and practical conservation solutions where we can intervene to help preserve coral biodiversity and genetic diversity. Decisive conservation actions are urgently needed to save our reefs.

Cryopreservation is a state-of-the-art tool that has been used successfully for decades to preserve biodiversity and genetic diversity in many wildlife species (Wildt 1992). The process works because, through a series of steps, water within the cell is extracted and replaced with a cryoprotectant or antifreeze. The partially dehydrated cell can then withstand the extraordinary stress of low temperature exposure, essentially entering a state of suspended animation (Mazur 1984). Cryopreservation can maintain the sample cold-but-alive for decades, thus offering much needed time to help resolve *in situ* conservation challenges. Cryopreservation is a maturing conservation tool that already has impressive milestones for coral. To date, the global community has cryopreserved coral sperm from over 50 species (https://nationalzoo.si.edu/center-for-species-survival/coral-species-cryopreserved-global-collaborators). These cryopreserved assets have been used to create new coral from these frozen sperm samples for restoration and assisted gene flow (Daly et al. 2022; Hagedorn et al. 2012; Hagedorn et al. 2017; Hagedorn et al. 2021) and proof-of-concept experiments have preserved coral larvae (Daly et al. 2018).

Coral reproduction occurs for most species only over a few days each year (Babcock et al. 1986; Bouwmeester et al. 2021). However, it is faltering in some areas of the world due to stressors, an important milestone in coral cryopreservation is to preserve small pieces of adult coral (0.5 cm^2^), also commonly called coral ‘microfragments’ (Koch et al. 2021; Page et al. 2018a) of a variety of species to help secure the genetic diversity and biodiversity of reefs. This type of cryopreservation would be relatively immune to climate change issues and could be accomplished throughout the year. However, before robust cryopreservation strategies for coral microfragments can be developed, basic cell sensitivities to chilling and cryoprotectant solutions must be tested, and the response of these fragments to these stressors must be monitored over time. Not only is it important to produce cryopreserved coral, but it is equally important to create a clear husbandry pathway to return these fragments to a land-based nursery setting post-thaw. This paper examined: 1) the sensitivity of the coral microfragments with their algal symbionts to chilling temperatures, given coral larvae and algal symbionts are known to be extremely sensitive to chilling (Hagedorn et al. 2006); 2) the response of the coral to complex cryoprotectant cocktails in terms of toxicity and how long it took them to start regrowing after this exposure; and, 3) methods to safely remove the algal symbionts from the coral fragment before cryopreservation resulting in bleached fragments, given the two symbiotic partners have very different membrane permeabilities to water and cryoprotectant (Hagedorn et al. 2009; Hagedorn et al. 2006). Methods for quantifying physiological responses to these stressors include confocal imaging, Pulse Amplitude Modulated fluorometry, imaging Pulse Amplitude Modulated fluorometry, light microscopy, standardized health metrics and bleaching color cards. A deep understanding of these types of detailed physiological stressors and metrics will be critical to help overcome the inevitable stress of cryopreserved coral fragments.

## Materials and methods

### Coral collection and Microfragmentation

*Porites compressa* colonies were collected from various reefs throughout Kāneʻohe Bay, Oʻahu, HI in accordance to our collecting permit from the Department of Land and Natural Resources from the State of Hawaiʻi (Special Activity Permit #2022-22 from the Hawaiʻi Institute of Marine Biology). A hammer and chisel were used to collect 10–15 cm portions from individual colonies. Care was taken to collect from colonies at different locations on each reef and throughout Kāneʻohe Bay to avoid collecting clones of the same genotype. Once collected, colonies were kept in outdoor aquaria with a filtered, flow-through seawater system at the Hawaiʻi Institute of Marine Biology on Moku o Loʻe.

Prospective colonies were progressively fragmented to yield uniformly sized microfragments (0.5 cm^2^) and glued (Super Glue Gel) onto a plastic sheet supported by a plexiglass plate (Page et al. 2018b). Microfragments were then allowed to heal for 2 weeks prior to experimentation.

### Solution Preparation

Given their size and complexity, future preservation of coral microfragments will likely require a cryopreservation process called vitrification, which avoids lethal ice formation by allowing the liquid within the system to enter a vitreous or glassy state. This process uses high concentrations of solutes (at very minimum what is being tested in this paper, and more frequently on the order of 7–10M) and ultra-rapid cooling and warming, at 10^4^ °C/min or higher rates of warming and cooling. The vitrification solutions were based off of previous methods and solutions used to cryopreserve coral larvae (Daly et al. 2018), but the cryoprotectants were reduced 20–24% and the trehalose was reduced by 23%. Two different strengths of the same vitrification solution (VS) were prepared for testing: (1) VS 80% (0.8 M dimethyl sulfoxide, propylene glycol and glycerol, 0.7 M trehalose in 0.3 M PBS (Phosphate Buffered Saline); and (2) VS 76% (0.75 M dimethyl sulfoxide, propylene glycol and glycerol, 0.7 M trehalose in 0.3 M PBS, see Table S1 for details).

### Pulse Amplitude Modulation Fluorometry

For these studies we used two types of Pulse Amplitude Modulated (PAM) fluorometer. The first was a Junior-PAM (Walz, Effeltrich, Germany) with a single fiber optic cable of approximately 1 mm in diameter for the toxicity and chilling experiments (Walz, Effeltrich, Germany). The second was an Imaging-PAM (IMAG-MAX/L; Walz, Effeltrich, Germany) for the bleaching experiments. Both PAM units measure functionality of Photosystem II in photosynthetic organisms. Specifically, photosynthetic yield was used to determine the approximate health and functionality of the algal symbiont after exposure to the various treatments (i.e., toxicity, chilling, bleaching).

For the toxicity and chilling experiments, three different points on each coral fragment were sampled at each of the 4–5 health assessment time points over time. For the bleaching experiments we used an Imaging-PAM, which is able to assess the photosynthetic yield across the entire microfragment. We chose to define five areas of interest (1 mm^2^ on each microfragment) with the Imaging-PAM, which allowed us to determine the photosynthetic yield of a much larger area of the microfragments. Generally, anything above a Photosynthetic Yield of 0.1, on a scale from 0 to 1, suggested functional symbionts.

### Chilling Sensitivity of Coral Fragments and their Algal Symbiont

During a vitrification experiment, the coral might experience a certain period of chilling. Cells or tissues that are extremely sensitive to chilling can experience ruptures in the cell membranes around 0 °C and might experience potential toxicity from this process. Therefore, we needed to explore the limit of what coral microfragments could tolerate and still recover from these exposures.

Preliminary experiments determined that coral microfragments could only withstand 1 min of chilling at 0 °C (see Supplementary Methods and Data, Fig. S1). Lower temperatures or longer exposures either led to immediate death or death within 2 weeks. Therefore, microfragments (n =11) were chilled at 0 °C for 1 min in cryovials with 1 ml FSW (0.22 µm filtered seawater) that had been pre-equilibrated to 0 °C for 10 min to determine how they would recover from this stress. After 1 min of chilling, microfragments were placed in 1 L of FSW at room temperature (∼ 22 °C) for 10 min and then were placed in individual 6-well plates with approximately 10 ml FSW. For the first 24 h of culture, microfragments were kept at 26 °C and 0 PAR and then given additional light (35–50 PAR) thereafter.

Chilled microfragments were assessed at 4 time points: 24, 72 h, 1-, and 2-wk post-chilling exposure. Each assessment included, 1) symbiont viability with a Junior PAM.; and 2) the integrity of the coral tissue with light microscopic imaging and health metric scoring (see Table 1 for details). After the 24 h assessment, microfragments were placed back into an incubator at 50 PAR and 26 °C. Coral fragments were cultured in 6-well plates where their plate and FSW were changed daily through the 72-h assessment, afterwards, they were moved to 5-L aquaria with running FSW through the 2-wk assessment at 26 °C and 35 PAR.

**Table 1.** Health metric scoring criteria for unbleached microfragments.

| **Score** | **Description of metric for unbleached microfragments** |
| --- | --- |
| **0** | Dead, coral tissue and algal symbionts released from skeleton, yellow/green color |
| **1** | Damaged polyp, green color, <10% intact coenosarc |
| **2** | Damaged polyp, pale, 25% intact coenosarc |
| **3** | Intact polyp, paling color, 50% intact coenosarc |
| **4** | Intact polyp, normal tissue color, 75% intact coenosarc |
| **5** | Intact polyp, normal tissue color, 100% intact coenosarc |

\*Higher score indicates greater health of the fragment

### Toxicity of Vitrification Solution to Coral Fragments

Coral fragments (n=11) were placed in vitrification solution of either VS76 or VS80 for 6 minutes in one step and then placed through a rehydration series (Table S2). The physiological effect of the solutions on the coral were not significantly different (Mann-Whitney test, p>0.05). Thus, the results from these two vitrification solutions were eventually pooled as one treatment, called VS. Coral microfragments were placed into individual 6-well plates with approximately 10 ml of FSW for recovery. Six-well plates were then placed in an incubator (26 °C), covered in aluminum foil (0 PAR) for the first 24 h post-cryoprotectant exposure, and 35–50 PAR, thereafter. Microfragments were assessed at 5 time points: 24 and 72 h, 1-, 2-, and 3-wk post-cryoprotectant exposure. Each assessment included: Junior PAM fluorescence reading, laser-scanning confocal microscopy imaging, light microscopy imaging and health metric scoring (Table 2). A health score of 0-5 was assigned to each coral fragment at each of the health assessment time points (24 and 72 h, 1 and 2 wk). The health score was based off of the following criteria: tissue loss and algal symbiont loss, tissue color, damaged or intact polyps, and intact coenosarc (Table 1).

**Table 2.** Health metric scoring criteria for bleached microfragments*.

| **Score** | **Description of metric for bleached microfragments** |
| --- | --- |
| **0** | Dead, tissue sloughing or >25% bacterial growth, >75% brown “freckles” or sheeting of dead/dying algal symbionts on surface of coral |
| **1** | >75% damaged polyps, <10% intact coenosarc, >50% brown “freckles” or sheeting of dead/dying algal symbionts on surface of coral |
| **2** | 50% damaged polyps, 25% intact coenosarc, >25% brown “freckles” or sheeting of dead/dying algal symbionts on surface of coral |
| **3** | >75% intact polyps, 50% intact coenosarc, <10% brown “freckles” or sheeting of dead/dying algal symbionts on surface of coral, no bacterial growth |
| **4** | Intact polyps, 75% intact coenosarc, <5% brown “freckles” or sheeting of dead/dying algal symbionts on surface of coral |
| **5** | Intact polyps, 100% intact coenosarc, no brown “freckles” or sheeting of dead/dying algal symbionts on surface of coral |

\*Higher score indicates greater health of the fragment

Coral fragments were kept in 6-well plates with FSW (changed daily) in the incubator through the 2-wk assessment, after that, they were placed in 5 L aquaria with running seawater through the 3-wk assessment.

Chilling and toxicity were not tested together, because during cryopreservation as the tissue starts to chill, the permeability of the cell membranes reduces the flow of cryoprotectant into the cells, thus limiting the impact of toxicity. Vitrification would be necessary for large complex tissues like coral microfragments. Generally, vitrification procedures involve dehydrating and equilibrating tissues in their vitrification medium at room temperature and then rapidly cooling the coral cryopreservation temperatures.

### Growth Response After Chilling and Cryoprotectants Exposure

After exposure to either chilling or vitrification solutions, coral fragments were cultured for two–three weeks, and then moved to a flowing seawater system whereby they were secured to plastic sheets supported by plexiglass plates and suspended in the flowing water. Every week, they were examined to determine whether they had recovered enough to put energy into new growth. We defined this by the production of one to two rows of coral polyps that form a full ring around the microfragment on the plastic sheet.

### Microfragment Bleaching

Preliminary experiments were done with vitrification of coral fragments and one concern was that the cryo-permeabilities of the algal symbionts and coral tissue to cryoprotectants were very different (1 h versus 3 mins, respectively), resulting in the symbionts dying and causing stress within the coral host. Trying to equilibrate the algal symbionts prior to cryopreservation resulted in the coral fragment dying. Therefore, we developed bleaching protocols that maintained the coral health with the caveat that we would be able to reintroduce the algal symbionts after thawing during the culture period. Three treatments were preliminarily assessed, including menthol (Wang et al. 2012), light, and menthol and light.

For subsequent bleaching treatments, the combination of menthol and light was used. Before the bleaching treatment, microfragments were imaged, assigned a health score (Table 2), and given a color rank assessed by Koʻa Card (Bahr et al. 2020) in order to determine any change in health during the bleaching process. During the day, microfragments were placed in aerated, 2 L aquaria with 0.58 mM menthol (99%, Sigma Aldrich, St. Louis, MO) in ethanol (190 proof, Decon Labs, Inc., King of Prussia, PA) in filtered seawater with aeration for approximately 6-7 h, 26 °C, ∼5 PAR. Then, they were transferred to a 26 °C incubator with 350 PAR for 17–18 h. The microfragments were cycled between the menthol bath and full light until they were fully bleached (i.e., had reached the lowest color ranking on the Koʻa Card). This took approximately 72 h–1 wk. At the end of the bleaching treatment, microfragments were imaged on a light microscope with a Lumenera Infinity 3s camera, assigned a health score (criteria based on Table 2), given a color rank assessed by Koʻa Card, imaged on a Zeiss LSM 710 laser-scanning confocal microscope, to determine the presence and viability of the algal symbionts was assessed by a Walz Imaging-PAM.

### Confocal Imaging of microfragments

Confocal imaging was used to assess coral viability of post-thaw microfragments (measuring GFP and algal symbionts) and to quantify the success of microfragment bleaching (mean fluorescent intensity of algal symbionts).

Each coral fragment was imaged using the Zeiss LSM 710 with a Zeiss Plan-Apochromat 5x/0.16 M27 objective. All coral fragments were imaged with the same acquisition settings: z-stack: 12 slices, range =330 um; image resolution: 2048 x 2048, (1700 um x 1700 um), 12-bit; pixel dwell: 1.57 microsec; pinhole size: 36 um, 1 AU. The image frame for all samples was entirely composed of coral coenosarc tissue and one polyp. No blank space occupied the image frame.

The excitation wavelength of 405 nm was applied using a Diode laser at 15% intensity. Two channels were created to capture the autofluorescence of the coral fragment. Channel 1 (515–575 nm) captured the autofluorescence of the coral host, Channel 2 (611-709 nm) captured the autofluorescence of the chlorophyll within the algal symbiont cells. The symbiont autofluorescence in the confocal image is used as a proxy for algal symbiont density within the coral tissue (Huffmyer et al. 2021).

For MFI analyses, Zen Black processing software (Zen 2.3 SP1 FP3 v.14.0.26.201) was used, all z-stack images were standardly formatted into maximum intensity projections. Fluorescence intensity data of the symbiont from the maximum intensity projections was pooled together within each experimental treatment (control and bleached) and averaged.

### Statistical Analyses

Measurements were represented by the means in all figures. All data were tested for normality and outliers using the ROUT method set at a sensitivity of 1%. For normally distributed paired data, parametric t-test were performed. If the data were not normally distributed, non-parametric tests (Mann-Whitney or Krukal-Wallis test) were done to test the differences amongst means. Where needed, Analyses of Variance (ANOVA) or Kruskal-Wallis tests were used to determine differences between groups (α=0.05). When groups were significantly different, posthoc tests were conducted using Dunn’s multiple comparisons tests. All error bars in the figures are represented by SEM. Statistical analyses were conducted in Prism 9.31 (GraphPad, San Diego, CA).

## Results

### Chilling and Toxicity Sensitivity of Coral Fragments and their Internal Symbiont

Preliminary experiments narrowed the chilling exposure time that the coral microfragments could tolerate (i.e., 0 °C for 0.5-, 1-, 2- and 4-min exposures were tested, see Supplementary Methods and Data, Fig S1). The coral tissue and their algal symbionts appeared to withstand chilling to 0 °C after 24 h for all exposure times except 4 min. However, survival/recovery was not clear until at least 1 wk after chilling exposure. The 2 and 4 min chilling exposure times caused corals to lose the majority of their tissue or die. This suggested that there were long-term consequences to low temperatures for the coral and that monitoring their overall physiology to 24 h post-thaw was not sufficient. The true measure of health and viability could only be measured approximately 1 to 2-wk post-treatment.

Therefore, for more detailed analyses, we examined the chilling sensitivity of the *P. compressa* microfragments at 0 °C for 1 min exposures over 2 weeks (Fig. 1). Untreated (control) microfragments maintained a uniform health metric of 5 throughout the treatment period (Kruskal-Wallis; *p>*0.05), and their mean photosynthetic yield did not vary greatly, although the initial readings at 0 and 24 h were lower than the later values over the two-week period in culture (Kruskal-Wallis *H*_(6)_=17.05; *p*<0.001; Dunn’s Multiple Comparison test). Compared to the control values at 0 h, the chilled microfragments’ health metrics (Fig. 1 a and b, blue bars) showed a decline followed by an improvement at 2-wk, with the 2-wk fragments comparable to control values (Kruskal-Wallis *H*_(5)_=17.33; *p*<0.001; Dunn’s Multiple Comparison test). During this recovery period, chilling caused a loss of coral and algal symbiont cells, which were observed surrounding the microfragments in the culture dishes for up to 72 h. These stressors caused the fragments to pale, before they recovered. At 72 h, the photosynthetic yield was lower than the control values, after which it returned to pre-treatment values (Kruskal-Wallis *H*_(5)_=17.09; *p*<0.004; Dunn’s Multiple Comparison test).

**Figure 1.**
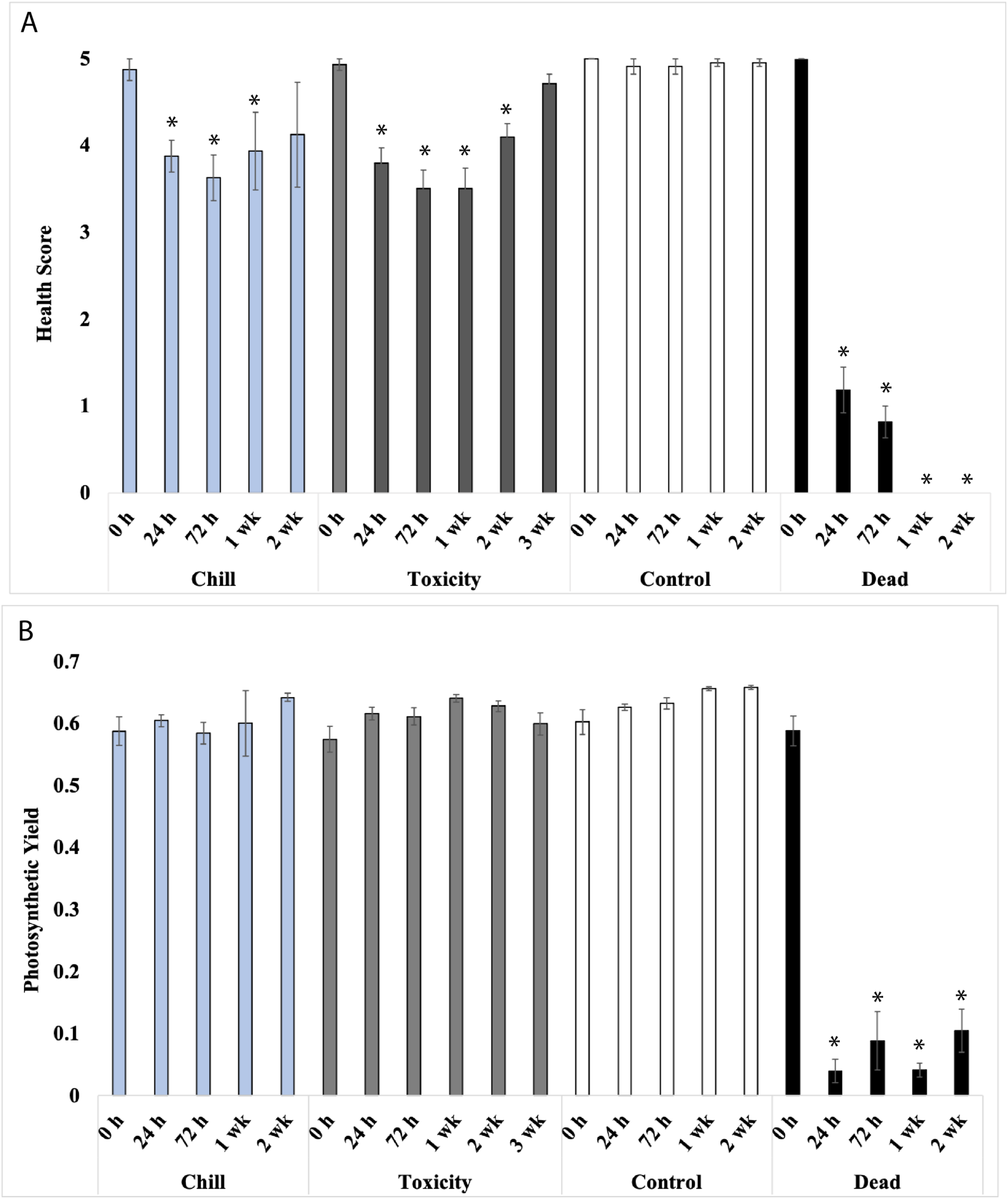
Physiological responses of *P. compressa* to chilling temperatures or toxic cryoprotectants. Microfragments from *P. compressa* were exposed to 1 min of chilling at 0 °C for 1 min (blue bars), or were exposed to two different vitrification media (black and gray bars) for 6 min at 22 °C then monitored for 2 to 3 weeks. **a.** Heath metric scores demonstrated a decrease in health, reflective of either loss of symbionts (during chilling) or a loss of coral cells during toxicity exposure followed by recovery in all cases after two to three weeks. **b.** The photosynthetic yield was relatively unchanged post-treatments although there were variations of the means of about 10% over time. Anything below a score of 2 on the health metric scale was not observed to recover. The * indicates a significant difference from the 0 h control health and photosynthetic yield values (*p*<0.05).

To determine how long microfragments could be exposed to the vitrification solution, preliminary toxicity experiments were conducted over several time points (1 to 5 min). Because the means were not different across exposure times (p> 0.05; Fig S2), we continued our experiments with a slightly longer exposure of 6 min and extended the time we monitored recovery from 2 wk to 3 wk.

Microfragments demonstrated some loss of coral cells and algal symbionts associated with some tissue retraction at the 24 h to 2 wk time-period reflecting poorer health, followed by a recovery at 3 wk (Fig. 1 a and b, gray bars; Kruskal-Wallis *H*_(6)_=57.08; *p*<0.0001; Dunn’s Multiple Comparison test.) There was no visible loss of algal symbionts observed during any cryoprotectant treatment, however, the photosynthetic yield was slightly lower at the 24 h, 72 h and 3 wk time points (but not at the 1 wk and 2 wk time points) when compared to the 1-wk control photosynthetic yields (Kruskal-Wallis *H*_(5)_=12.21; *p* =0.0159; Dunn’s Multiple Comparison test). These experiments were critical to understand the physiological changes that the coral might undergo prior to cryopreservation, during which they will be exposed to low temperature stressors, as well as toxicity.

### Growth Response After Chilling and Cryoprotectant Exposure

After the coral microfragments were exposed to chilling or toxicity, they were kept in recovery for two-three weeks in the lab and then returned to running seawater tanks to determine how long it would take them to recover growing. Not all fragments were followed through re-growth, the sample size for each treatment was as follows: 1) no treatment (n=9), 2) chilled to 0 °C for 1 min (n=9), or 3) exposed to VS (n=7). The mean time for each group to begin growing in our seawater system was two months and there was no difference between any of the treatments (Kruskal-Wallis test; *p*>0.05), suggesting that once the microfragments had recovered in the laboratory at 2–3 weeks, their previous treatment did not affect their future growth. Specifically, the timing for re-growth was Control =60.0 ± 6.3; Chilling =65 ± 5.1; Toxicity =56.1 ± 5.0 days.

### Confocal Imaging

In this study, confocal imaging was used to try to understand the patterns of the green fluorescent protein (GFP) and algal symbionts in the tissues of both live and dead coral fragments, as well as to quantify the success of microfragment bleaching.

Living coral fragments have a well-defined distribution of the auto-fluorescent green fluorescent protein (GFP), and the discrete distribution of their auto-fluorescent algal symbionts, which generally surround the tentacles and polyp mouth. However, each genotype has a unique distribution of these fluorescent signatures that define these living fragments. The control pattern of fluorescence for live and dead is shown in Fig 2. Dead coral have a different fluorescent pattern. After a coral fragment has gone through several freeze-thaw cycles, ice crystals disrupt their membranes, and the fragments die. This causes the GFP and algal symbiont fluorescent signals to become disaggregated and disorganized, producing a smeared appearance, although the GFP signal remains up to 72 h (Fig. 2). In fact, when the mean fluorescence intensity of the live and dead corals was compared, there was no difference in GFP or algal symbiont fluorescence. Because of the longevity of the GFP in the tissue, the presence of this signal was not deemed a good indicator and could not be used to quantify viability of post-thaw coral fragments.

**Figure 2.**
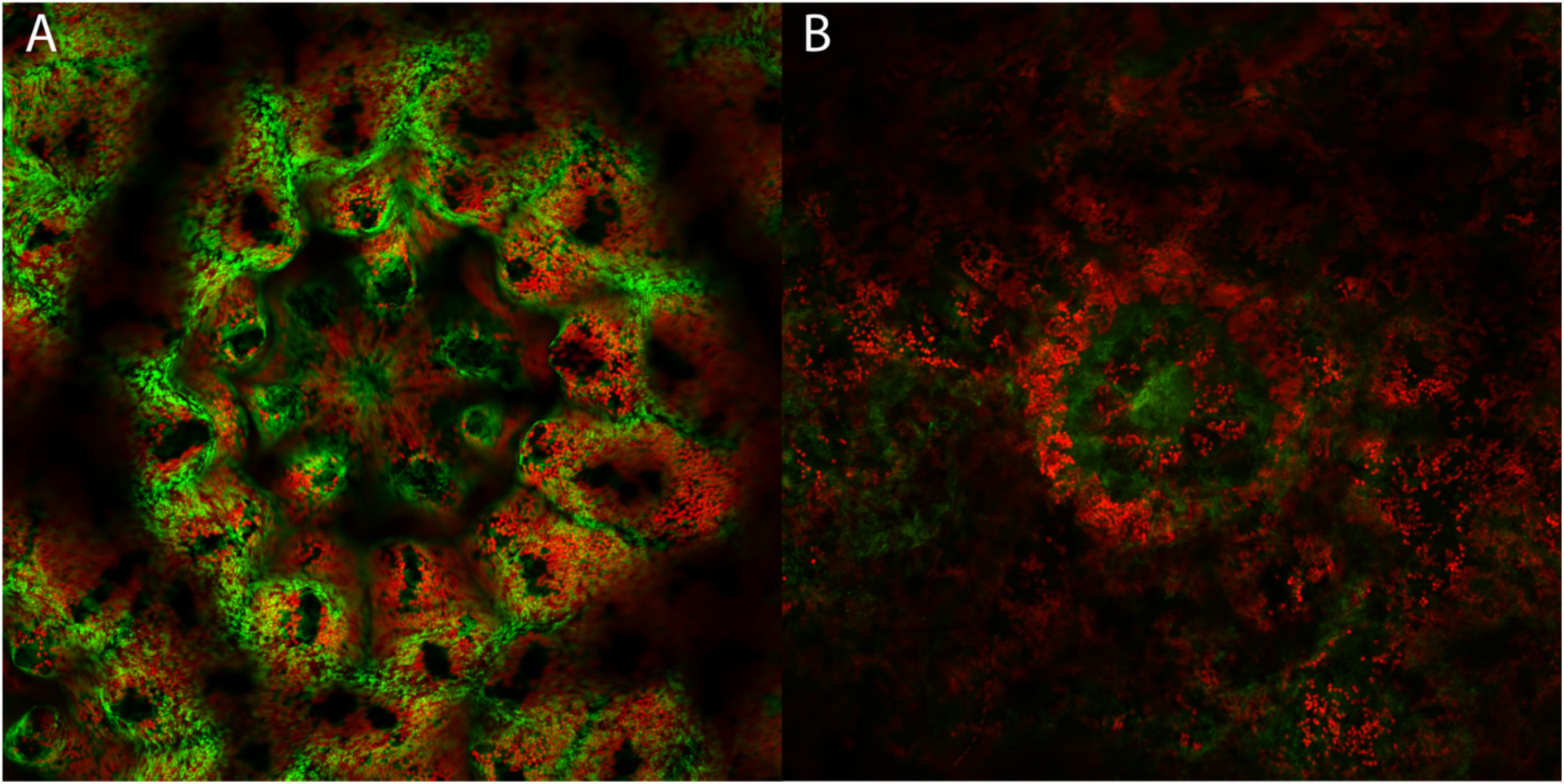
Confocal images of a polyp from a Live and Dead coral fragment after 24 h in culture. **A.** Live polyp, 24 h: Merged image of the auto-fluorescent symbiotic algae (red) and the auto-fluorescent green fluorescent protein of the coral (GFP, green). Note how tightly organized the GFP and symbionts are around the polyp mouth and tentacles. The confocal image clearly shows the morphology of the polyp skeleton and tentacles. **B.** Dead polyp, 24 h: Merged image of the auto-fluorescent symbiotic algae (red) and the auto-fluorescent GFP of the coral (green). Note the disorganized pattern of the GFP and symbionts fluorescence around the polyp and tentacles. The polyp skeleton and tentacles were degraded and appear blurred. The symbiont fluorescence is scattered across the image and the GFP is blurred.

Additionally, confocal imaging was used to determine the success of intentional microfragment bleaching to assess whether the symbionts disappear from the tissue or were non-functional. Preliminary experiments determined that a combined menthol and light bleaching treatment resulted in an 83% decrease in Mean Fluorescence Intensity between the wavelengths fluoresced by the algal symbiont after 72 h of exposure (Fig. 3). Specifically, in this image, the control had a Mean Fluorescence Intensity=284.6; light treated=78.2; menthol=89.1, and menthol and light=42.1. Additionally, all treatments maintained a health metric score of 5, therefore, the menthol and light bleaching treatment was used for subsequent assessments.

**Figure 3.**
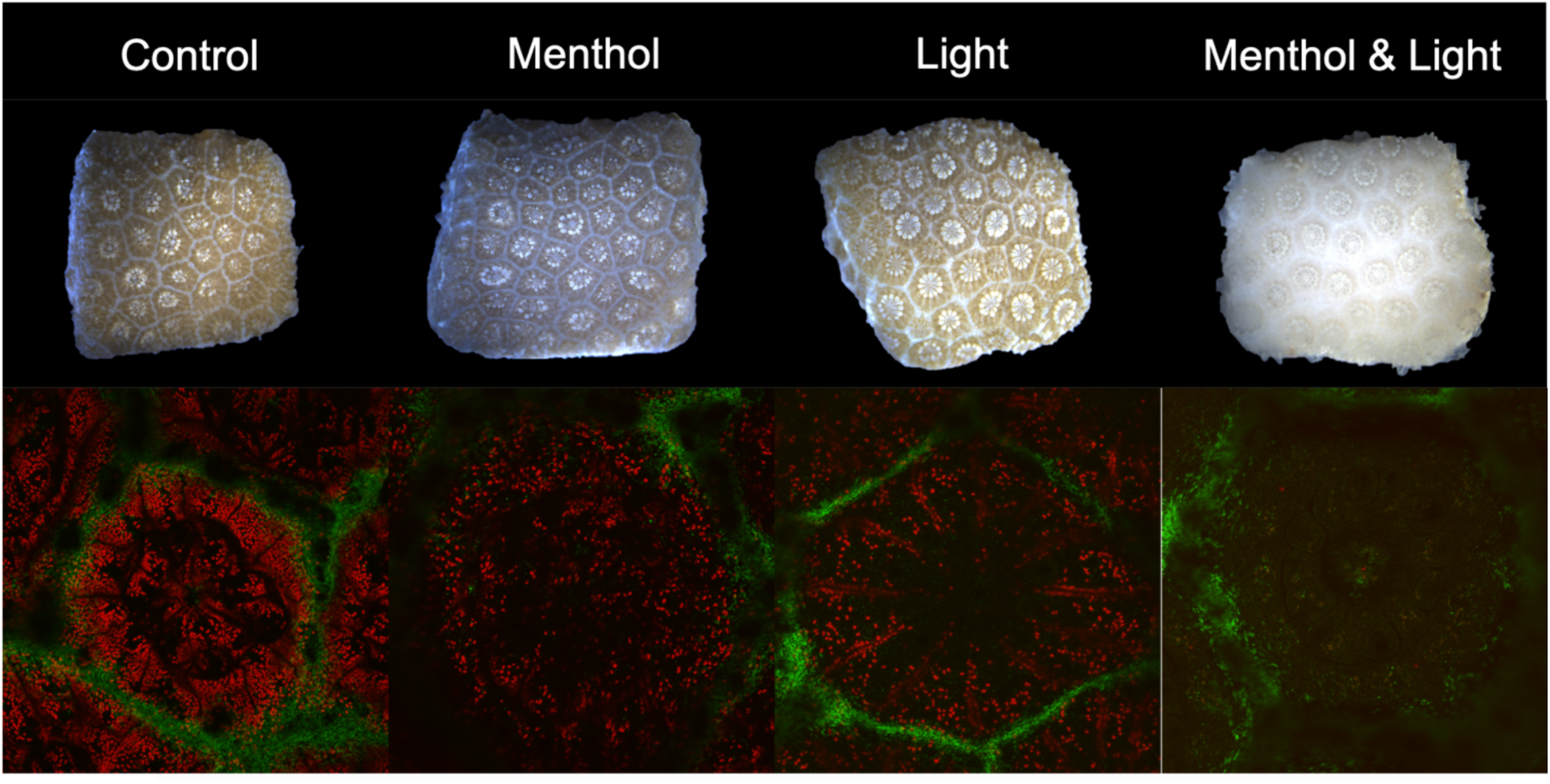
Coral fragments were treated for 3 days with one of the bleaching treatments and the density of symbionts were assessed. **Upper panel**: Light images of coral fragments that were ‘bleached’ using Menthol, Light, and Menthol & Light. **Lower panel**: Confocal images of the same coral fragments but at the polyp level. The merged images layer the GFP (green) and autofluorescence of the algal symbionts (red) into one image. Preliminary data show that the combination of Light & Menthol reduced the density of symbionts the most.

In a more detailed study, we used the distribution of the algal symbionts throughout a z-stack to determine Mean Fluorescence Intensity. Menthol and light bleached microfragments (n =25), were imaged with confocal microscopy, Imaging-PAM, and given a health metric score to determine whether this treatment would significantly reduce the algal symbiont population without seriously compromising the health of the coral animal. Coral microfragments were bleached for 72 h–1 week and the loss of their symbionts was monitored. Paired microfragments from the same genotypes (n =25) were left untreated or bleached with menthol and light (Fig. 4). These pairs demonstrated a 78% loss in the Mean fluorescence Intensity or the number of symbiont-like fluorescing particles (Controls =212.1 ± 19.6; Bleached =47.0 ± 2.5; two-tailed t-test T(24) =8.90, p<0.0001).

**Figure 4.**
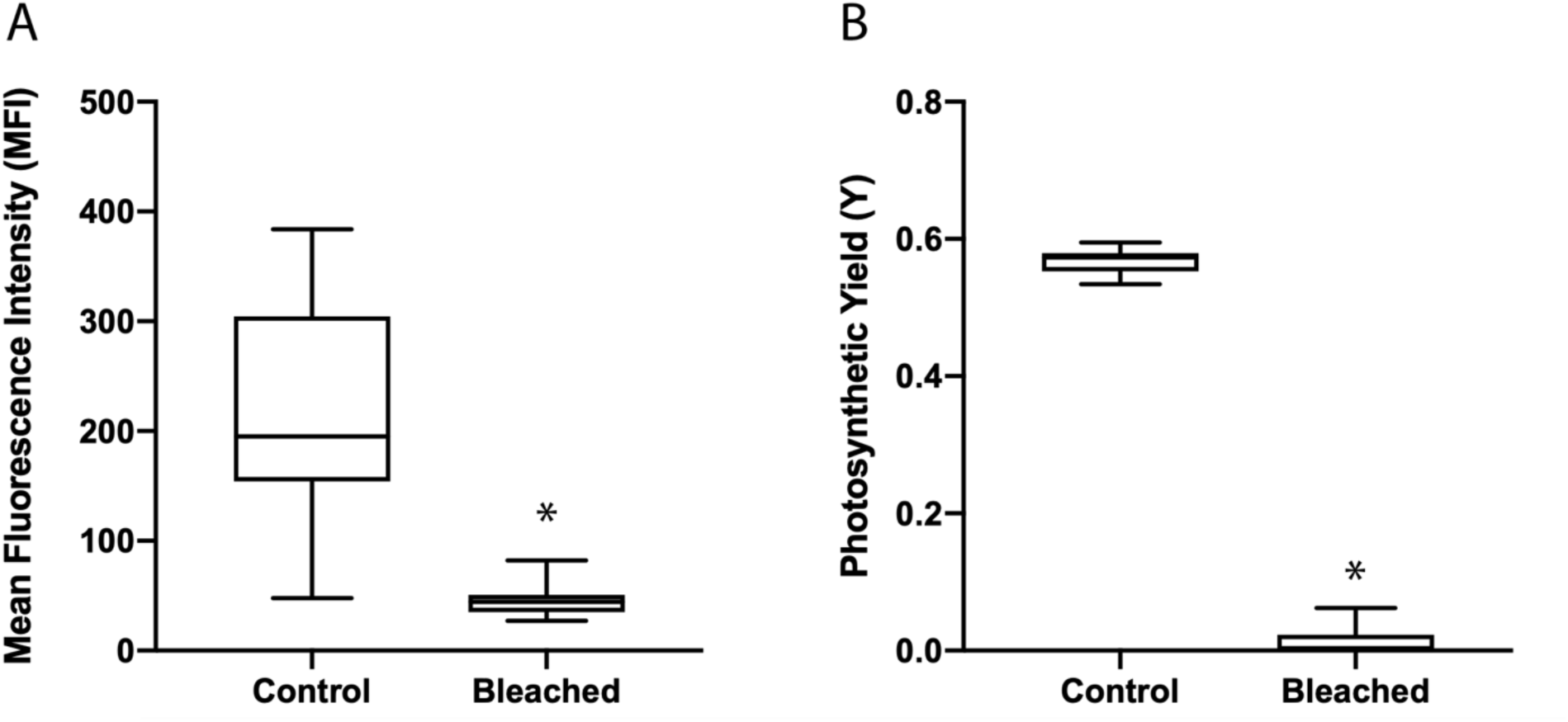
Coral microfragments were bleached up to 1 week and the loss of their symbionts was monitored with confocal microscopy and Imaging-PAM. **a)** Paired microfragments from the same genotypes (n=25) were left untreated or bleached with menthol & light. These pairs demonstrated a 78% loss in in the Mean fluorescence Intensity or the number of symbiont-like fluorescing particles (Controls=212.1±19.6; Bleached=47.0±2.5**). b)** When bleached and untreated microfragments from the same genotypes (n=10) were examined with an Imaging-PAM, a 98% loss in Photosynthetic Yield (Y) was observed. The control fragments had a mean Y-value of 0.567±0.006, whereas the bleached values were reduced to 0.013±0.007, suggesting that none of the remaining symbiont-like particles in the bleached fragments were functional. Means with * were different *p*<0.001, paired parametric t-test (a) and nonparametric non-paired Mann-Whitney U test (b), all errors represented by SEM.

When paired, bleached and untreated microfragments from the same genotypes (n=10) were examined with an Imaging-PAM, a 98% loss in photosynthetic yield (Y) was observed. The control fragments had a mean photosynthetic yield value of 0.567 ± 0.006, whereas the bleached values were reduced to 0.013 ± 0.007 (Wilcoxon matched-pairs test, p=0.0039), suggesting that none of the remaining symbiont-like particles observed in the bleached fragments with confocal microscopy were functional.

Each fragment was assessed under a dissecting microscope (Olympus BX41, magnification 20x) and given a score based on the rubric in Table 1 (control/unbleached) or Table 2 (bleached). Thus, the combined use of menthol and light reduced symbiont distribution and function throughout the coral tissue and maintained good health metric scores, 4.0 ± 0.3 (Table 3). There appeared to be a seasonal difference observed in the health of the bleached fragments. This difference was not statistically significant, however, it was observed that coral fragments that were bleached during the winter maintained higher average health scores (4.7 ± 0.2) than fragments that were bleached during the summer (3.1 ± 0.4).

**Table 3.** Differences in the health metric data for menthol and light bleached microfragments (bleaching over 72 h) observed seasonally.

|  | Winter | Summer | 72 hr<br>(combined) |
| --- | --- | --- | --- |
| **Average Health Score**<br>**(SEM)** | 4.7 ± 0.2 | 3.1 ± 0.4 | 4.0 ± 0.3 |
| **# of Genotypes** | 6 | 8 | 13 |

## Discussion

The goal of this paper was to understand and assess the physiological stressors that might underlie coral microfragment cryopreservation and the long-term consequences of these physiological exposures to continued coral growth in a land-based nursery. Coral microfragments initially responded negatively to chilling and toxic conditions, but recovered over a three-wk period in the laboratory. Once the microfragments had recovered in the laboratory and were placed in running seawater, they began to grow within 2 months, and their previous treatment did not affect their future growth. This may indicate that cryopreservation will cause some short-term stress, will recover and will not cause any long-term physiological impacts for the coral.

It was expected that the GFP intensity would wane more quickly over time in the dead fragments, so that they might be used as a post-thaw indicator of viability. However, the confocal imaging demonstrated that this was not the case up to 72 h, instead the pattern of GFP in and around the coral polyp changed from a very distinct roseate pattern to a more diffuse, distributed pattern. In contrast, warmed and cooled coral (±5°C) did demonstrate a loss of symbionts and GFP concentration (Roth & Deheyn 2013), so the persistence of the GFP signal in the dead coral was surprising.

A promising avenue to successfully cryopreserved something as large as a coral fragment is a technique called isochoric vitrification. Most biological matter is cooled under constant pressure or isobaric conditions at atmospheric pressure, which allows the sample to change volume. Moreover, only samples of relatively small size (∼100 µm in diameter), such as a human embryo, can be easily vitrified with these methods. However, emerging techniques aim to preserve biological material at constant volume (Rubinsky et al. 2005), confining the system and denying it access to the atmospheric pressure reservoir. This isochoric cryopreservation processes generally employs only a single step – cooling whereby the system is in a perfectly confined, constant-volume chamber. The technology does not require moving parts, mechanical work, and can be used on much larger samples. Therefore, iusochoric vitrification may be ideal for processes involved in field cryopreservation of coral microfragments near or on reefs.

To date, three general categories of isochoric cryopreservation have been developed, each inhabiting different thermodynamic regimes. The first is isochoric supercooling, which operates at a metastable local equilibrium, holds the liquid within the system in a supercooled state absent ice at temperatures ∼0–20°C (Consiglio et al. 2022; Powell-Palm et al. 2021) The second is isochoric freezing, which operates at global thermodynamic equilibrium, involves the partial freezing and self-pressurization of the system at temperatures between approximately 0 and −30°C (Powell-Palm et al. 2020; Wan et al. 2018). Finally, the third is isochoric vitrification (Zhang et al. 2018), which operates at a far-from-equilibrium steady state. This involves cooling the system rapidly to temperatures beneath the glass transition of the contained liquids (typically using liquid nitrogen submersion), almost completely arresting biochemical and transport processes while avoiding deleterious ice crystallization.

Given the demonstrated sensitivity of coral microfragments to mild chilling temperatures, this study suggests that isochoric vitrification may present a suitable technique to successfully cryopreserve them. How might this new vitrification process work for coral microfragments? According to this study, corals could withstand chilling temperatures up to 1 min, and complex vitrification cocktails for up to 6 min. Using isochoric vitrification, small volumes of vitrification medium (∼5 ml) can be frozen to liquid nitrogen temperatures (−196 °C) in less than 2 min. The amount of time where the microfragments might hover at chilling temperatures (0 to −10 °C) either on cooling or warming is less than 20 sec, well below the one-minute threshold found for chilling. Furthermore, previous work (Zhang et al. 2018) has found that isochoric confinement can reduce the CPA concentrations required to vitrify a given solution (as compared to conventional isobaric vitrification), which may enable use of minimally toxic cryoprotective solutions that would otherwise be susceptible to destructive ice formation.

There are calls for interventions to help restore coral reefs (National Academies of Sciences & Medicine 2019) and coral cryopreservation of all types is a maturing tool to aid in these conservation actions. However, engineers, coral biologists and cryobiologists must partner to help develop the tools to cryopreserve larger and larger cells and tissues and take the field into the future. One such endeavor, supported by the US National Science Foundation, is called APT-Bio, an Engineering Research Center (ERC) for Advanced Technologies for the Preservation of Biological Systems (https://www.atp-bio.org/). As part of this ERC, there is an emphasis on developing new medical technology to cryopreserve human organs. In terms of complexity of the tissues, the differences in the cryo-permeabilities of coral tissue and coral symbionts, the cryopreservation of coral microfragments might be considered almost as complex as human organs, such as an embryonic kidney or heart. Toward this end, there is emerging technology that may be a good candidate for coral. Restoration processes might benefit from the development of coral microfragment cryopreservation by allowing the safe preservation and reanimation of hundreds of thousands of small fragments potentially encompassing many of the coral species in the wild. Moreover, because there is such a small footprint for these frozen assets (compared to live assets in captivity), sufficient biodiversity can be maintained within a population to ensure robust repopulation efforts.

Some recent models suggest that time for these types of intervention processes is growing short (Dixon et al. 2022; Kalmus et al. 2022). If the cryopreservation of coral fragments is to be successful, there remain many unanswered questions about how many microfragments must be preserved, and when and where they might be collected from. However, this should not stop the scientific community from moving forward as quickly as possible to develop the technology fully, train professionals and bank the biodiversity in our oceans while it is still remains. Novel *ex situ* conservation strategies, such as genetic biorepositories holding cryopreserved coral fragments, hold strong promise to help offset many of the anthropogenic threats facing coral reefs today.

### Statements and Declarations

The authors have no relevant financial or non-financial interests to disclose.

## Acknowledgements

The authors would like to thank Dr. Shayle Matsuda for his advice and guidance on bleaching coral. We would like to thank the Coral Resilience Laboratory for the generous use of their Imaging-PAM. We would like to thank our summer interns: Kendall Fitzgerald and Morgan Brooks for their assistance in microfragment husbandry. HIMB contribution [# xxx] number will be added before publication.

## Ethics Declarations

On behalf of all authors, the corresponding author states that there is no conflict of interest.

## Author Contributions

Conceptualized and Executed Experiments: MH, CL, RP, JD, MM, MPP, JC;

Data Analysis: JB, CL, RP, MH, AC;

Husbandry: CP, RP, CL;

Writing and Reviewing Paper: JB, JD, CL, RP, CP, MH, MM, MPP, AC

