## Supplemental methods and figures for "Metrics of Coral Microfragment Viability"

### Supplementary Methods and Data

*Preliminary Chilling Experiments*

Coral microfragments (n=10 genotypes) were exposed to a chilling temperature of 0 °C to determine how long a coral fragment could experience chilling before it had an adverse effect on them. The fragments were held at the chilling temperatures for 0 sec, 30 sec, 1, 2 and 4 min, and then returned to room temperature post-treatment. The health metrics of the coral and the photosynthetic yield (a measure of symbiont health) were monitored over 1 wk.

The coral tissue and their symbionts withstood chilling to 0 °C at 24 h well, however, at 1 wk, most of the coral tissue and their symbionts were dead in the longer time exposures (2 and 4 min). This suggested that there were long-term consequences to low temperatures for the coral and that monitoring their overall physiology to 24 h post-thaw was not sufficient. The true measure of health and viability could only be measured approximately 1-wk post-treatment (see Fig. S1 a and b).

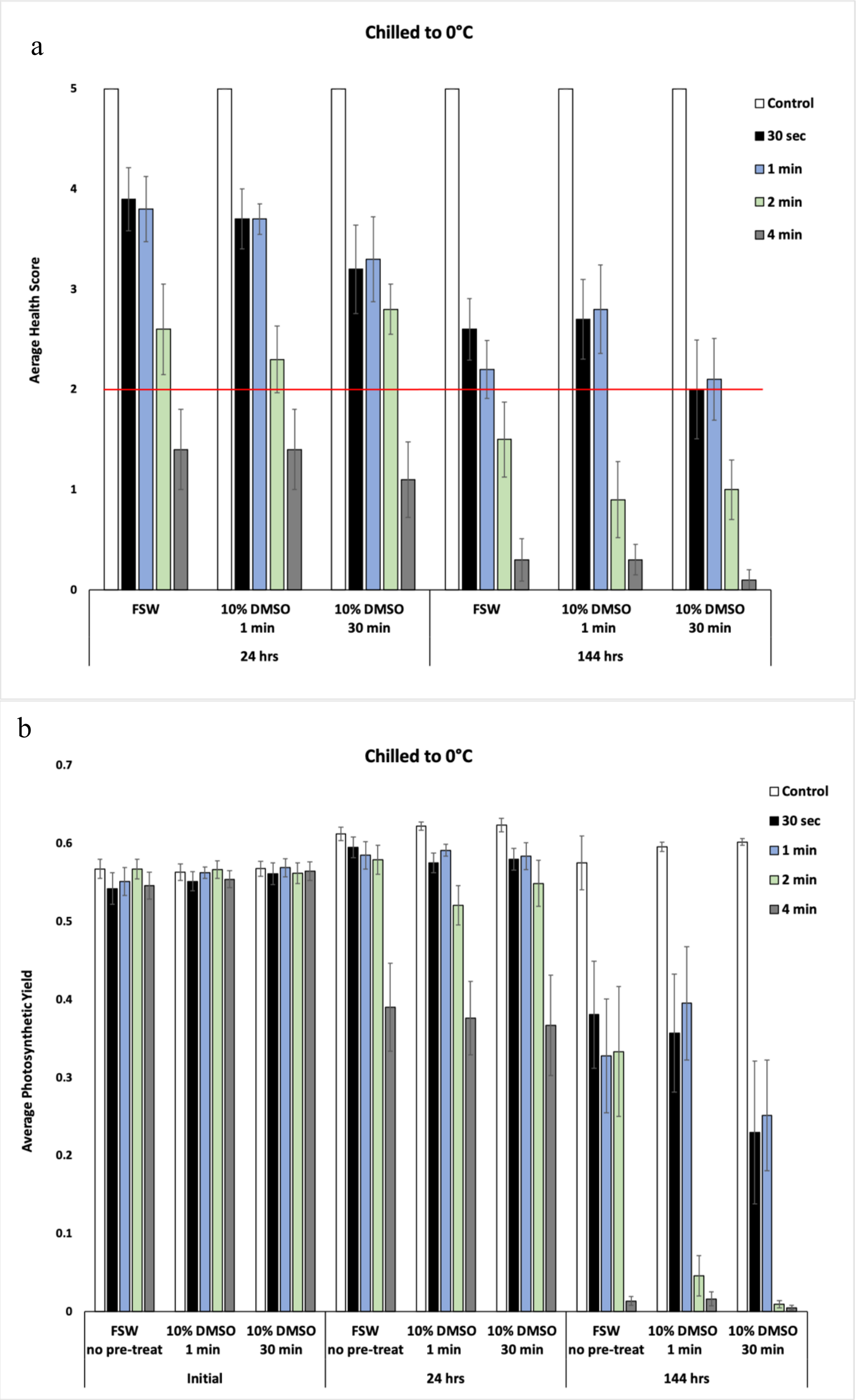


**Figure S1** The sensitivity of coral tissue (a) and their symbionts (b) to chilling temperatures. For the coral tissue, the red line in (a) indicated the health metric above which coral are most likely to survive. At 1 min or less of chilling at 0°c, there was some survival of coral tissue and their symbionts, but times beyond that there was none (a).

*Preliminary Toxicity Experiments*

Coral fragments (n=10 genotypes) were exposed to the vitrification solution (VS76) at 21 °C for 0, 1, 3 and 5 min, then moved back into filtered seawater and held in 6-well plates in an incubator over two wk. 6-well plates and seawater were changed daily. The health metrics of the coral was monitored over 2 wk.

The toxicity of the vitrification solution (VS76) over the time tested (1 to 5 min) did not impact the mean survival of the coral fragments over time ($p > 0.05$; Fig S2).

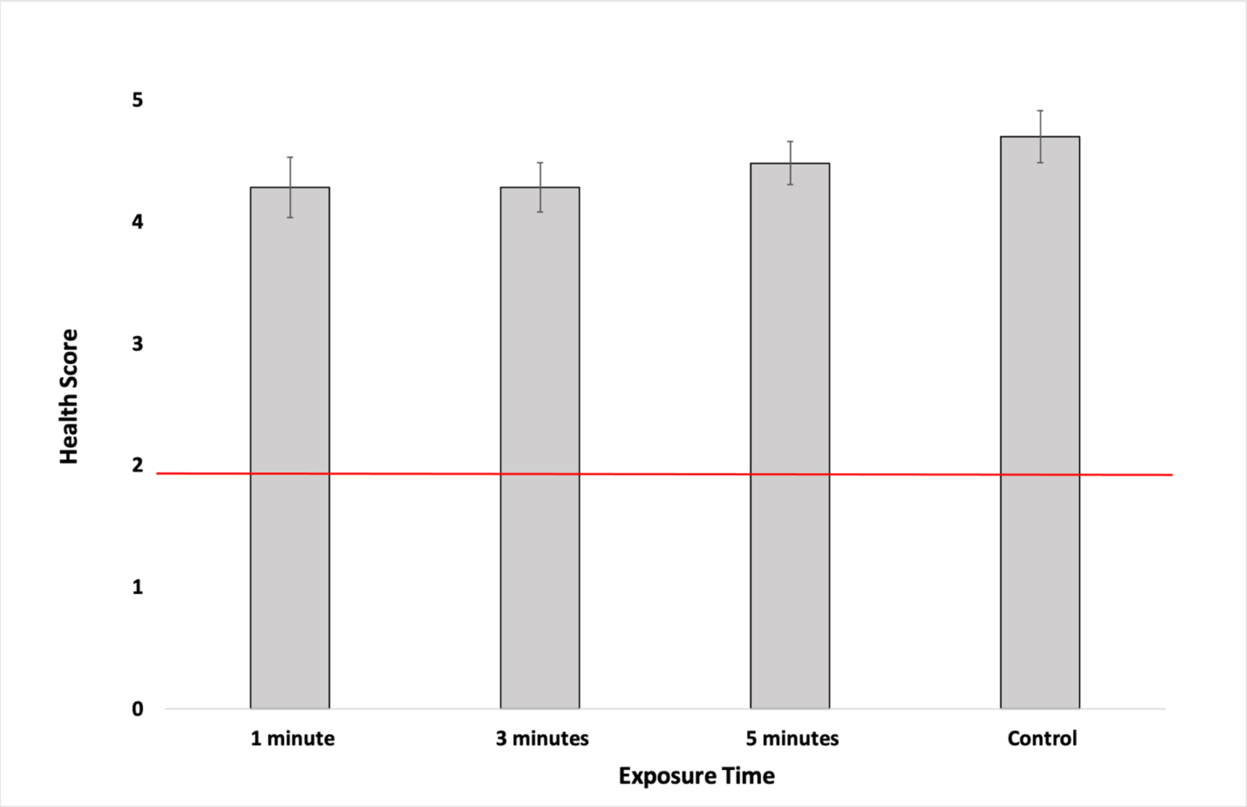


**Figure S2** Toxicity of the vitrification solution to the coral fragments over 2 weeks. The red line indicated the health metric above which coral are most likely to survive. There was no difference in the survival of the coral fragments to two weeks based on the length of exposure to the vitrification medium ($p > 0.05$, ANOVA)

Table S1 Description of the vitrification solutions used for toxicity experiments. Note the VS76 had the identical cryoprotectant concentrations, however it had ~13% more solute.

| **Component** | **Molar Mass** | **VS76** | **VS80** |
|---|---|---|---|
| **Phosphate buffer saline in deionized water (**Gibco 10x concentration; 70011-044 500 ml) | 18.1 | 35.4 g | 30.77 g |
| **Dimethyl sulfoxide (**Sigma Aldrich, D8418-500ML) | 78.13 | 2.75 g | 2.75 g |
| **Propanediol** (Sigma Aldrich; 398039-500ML) | 76.01 | 2.68 g | 2.68 g |
| **Glycerol** (Sigma Aldrich; G7893-500ML) | 92.09 | 3.24 g | 3.24 g |
| **Trehalose (**Swanson; SWU478) | 342.3 | 10.57 g | 10.57 g |

Table S2 Description of the rehydration series for the coral microfragments. FSW represents 0.22 µm filtered seawater

| **ACTION** | **Time in solution** |
|---|---|
| 1 ml 50% Vitrification Solution in FSW | 30 sec |
| Add 1.8 ml FSW to dish, gently pipette to mix | 30 sec |
| Add 3 ml FSW to dish, gently pipette to mix | 1 min |
| Add 9 ml FSW, gently pipette to mix | 1 min |
| Add 18 ml FSW, gently pipette to mix | 1 min |
| Flood the dish with 20 ml FSW | 1 min |
